# Alignment-based protein mutational landscape prediction: doing more with less

**DOI:** 10.1101/2022.12.13.520259

**Authors:** Marina Abakarova, Ćeline Marquet, Michael Rera, Burkhard Rost, Elodie Laine

**Affiliations:** Sorbonne Université, CNRS, IBPS, Laboratory of Computational and Quantitative Biology (LCQB), UMR 7238, Paris, 75005, France; Université Paris Cité, INSERM UMR U1284, 75004 Paris, France; Department of Informatics, Bioinformatics and Computational Biology - i12, TUM-Technical University of Munich, Boltzmannstr. 3, Garching, 85748 Munich, Germany; TUM Graduate School, Center of Doctoral Studies in Informatics and its Applications (CeDoSIA), Boltzmannstr. 11, 85748 Garching, Germany; Institute for Advanced Study (TUM-IAS), Lichtenbergstr. 2a, Garching, 85748 Munich, Germany; TUM School of Life Sciences Weihenstephan (TUM-WZW), Alte Akademie 8, Freising, Germany; Institut universitaire de France (IUF)

**Keywords:** genotype-phenotype relationship, protein mutation, multiple sequence alignment, deep mutational scan, evolution

## Abstract

The wealth of genomic data has boosted the development of computational methods predicting the phenotypic outcomes of missense variants. The most accurate ones exploit multiple sequence alignments, which can be costly to generate. Recent efforts for democratizing protein structure prediction have overcome this bottleneck by leveraging the fast homology search of MMseqs2. Here, we show the usefulness of this strategy for mutational outcome prediction through a large-scale assessment of 1.5M missense variants across 72 protein families. Our study demonstrates the feasibility of producing alignment-based mutational landscape predictions that are both high-quality and compute-efficient for entire proteomes. We provide the community with the whole human proteome mutational landscape and simplified access to our predictive pipeline.

**Significant statement:** Understanding the implications of DNA alterations, particularly missense variants, on our health is paramount. This study introduces a faster and more efficient approach to predict these effects, harnessing vast genomic data resources. The speed-up is possible by establishing that resource-saving multiple sequence alignments suffice even as input to a method fitting few parameters given the alignment. Our results opens the door to discovering how tiny changes in our genes can impact our health. They provide valuable insights into the genotype-phenotype relationship that could lead to new treatments for genetic diseases.

## Introduction

In recent years, tremendous progress has been achieved in the prediction of protein 3D structures and mutational landscapes (com, 2022; Laine et al., 2021) by leveraging the wealth of publicly available natural protein sequence data (Mirdita et al., 2022; Delmont et al., 2022; uni, 2023; Jumper et al., 2021; Nayfach et al., 2021; Camarillo-Guerrero et al., 2021; Mitchell et al., 2020; Levy Karin et al., 2020; Steinegger and Söding, 2018; Suzek et al., 2015; Nordberg et al., 2014). State-of-the-art predictors capture arbitrary range dependencies between amino acid residues by implicitly accounting for global sequence contexts or explicitly exploiting structured information coming from alignments of evolutionary related protein sequences. Very efficient algorithms, *e.g.* MMseqs2 (Steinegger and Söding, 2017), allow for identifying homologous sequences and aligning them on a mass scale. Others relying on profile hidden Markov models (HMMs), such as JackHMMer/HMMer (Eddy, 2011), carefully generate very large families, achieving a very high sensitivity. Several large-scale resources like Pfam (Mistry et al., 2021) and ProteinNet (AlQuraishi, 2019) give access to pre-computed multiple sequence alignments (MSAs) built from profile HMMs. These MSAs are associated with curated protein families in Pfam, or with experimentally resolved protein 3D structures in ProteinNet. The depth, quality, and computational cost of a MSA are important factors contributing to its effective usefulness. Nevertheless, precisely assessing the impact of expanding or filtering out sequences on predictive performance is difficult. For protein structure prediction, Mirdita and co-authors showed that AlphaFold2 original performance could be attained with much smaller and cheaper alignments through the MMseqs2 (Steinegger and Söding, 2017)-based strategy implemented in ColabFold (Mirdita et al., 2022). This advance makes accurate protein structure prediction more accessible and applicable at a much larger scale.

In this work, we aimed at testing whether the same gain could be obtained for mutational outcome prediction. We compared the prediction accuracy achieved by Global Epistatic Model for predicting Mutational Effects (GEMME) (Laine et al., 2019) from MSAs generated using the ColabFold’s MMseqs2-based protocol (Mirdita et al., 2022; Steinegger and Söding, 2017) versus three workflows relying on profile HMMs (AlQuraishi, 2019; Mistry et al., 2021; Notin et al., 2022) (**Figure 1**). GEMME is a fast unsupervised MSA-based mutational outcome predictor relying on a few biologically meaningful and interpretable parameters. It performs on-par with statistical inference-based methods estimating pairwise couplings (Hopf et al., 2017) and also deep learning-based methods, including family-specific models (Frazer et al., 2021; Shin et al., 2021; Trinquier et al., 2021; Riesselman et al., 2018) as well as high-capacity protein language models trained across protein families (Notin et al., 2022; Marquet et al., 2021; Meier et al., 2021) (see also (Trinquier et al., 2021; Marquet et al., 2021; Laine et al., 2019) for quantitative comparisons). GEMME is freely available for the community through a stand-alone package and a web server. It proved useful for discovering functionally important sites in proteins (Tsuboyama et al., 2023; Cagiada et al., 2023), classifying variants of the human glucokinase (Gersing et al., 2023) and transmembrane proteins (Tiemann et al., 2023), among others, and for deciphering the molecular mechanisms underlying diseases such as the Lynch syndrome (Abildgaard et al., 2023).

**Fig. 1:**
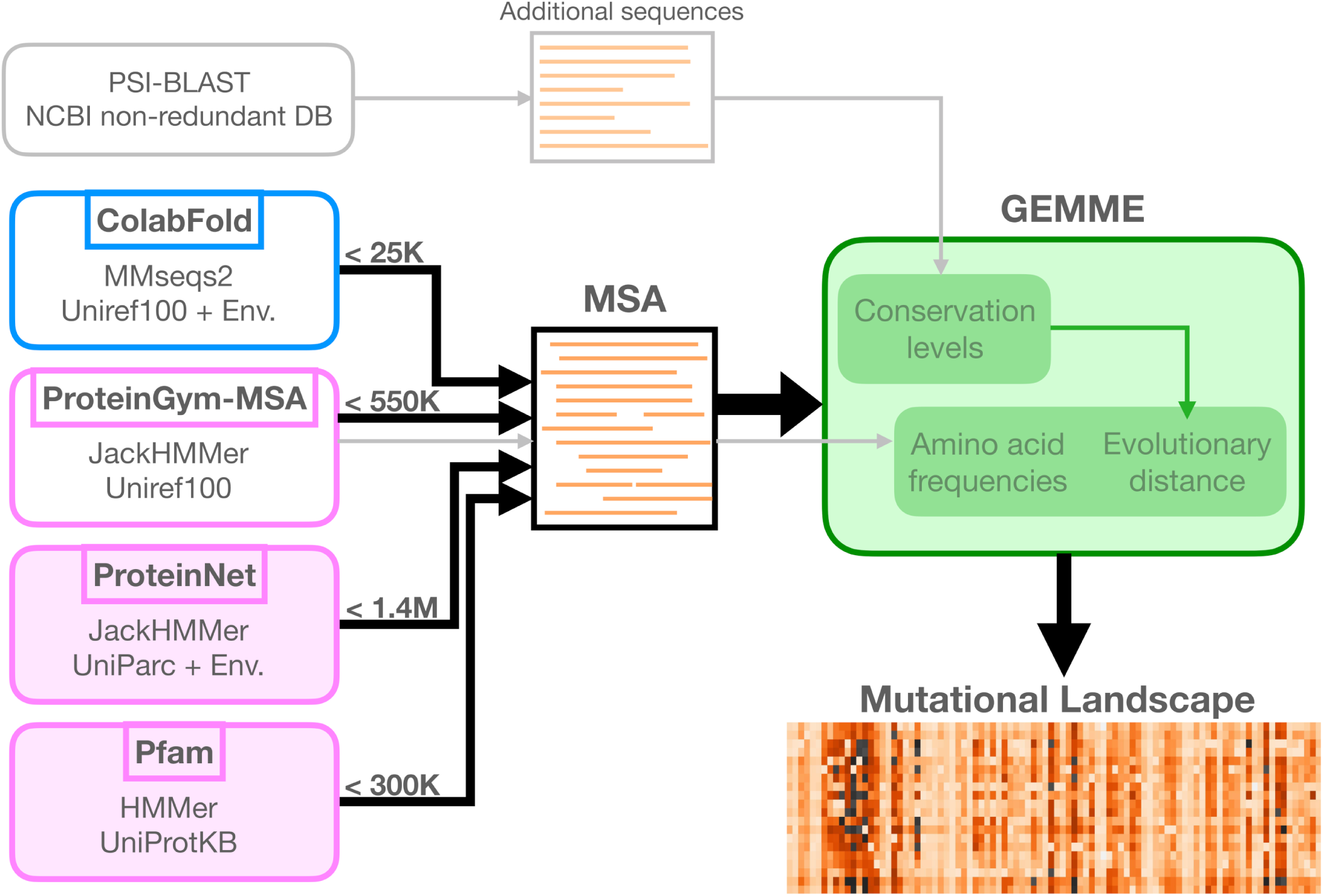
Schematic representation of the workflow. GEMME computes and combines conservations levels, amino acid frequencies and evolutionary distances to predict protein mutational landscapes. The original protocol (Laine et al., 2019), illustrated with grey arrows, used PSI-BLAST against NCBI’s non-redundant database to infer conservation levels, and additionally exploited an input MSA generated with JackHMMer against UniRef100 to compute amino acid frequencies and evolutionary distances. In the present work, GEMME computed all measures from a single input MSA (see black arrows). We assessed four MSA generation protocols and resources, one relying on a many-to-many sequence search (in blue) and the three others relying on profile HMMs (in pink). Two resources (filled rectangles) provide large amounts of MSAs, covering virtually all protein families or all proteins with an experimentally resolved 3D structure. For each protocol or resource, we indicate the maximum number of sequences in the considered MSAs, ranging from 25 thousands to 1.4 millions.

As GEMME optimized only a few free parameters (Laine et al., 2019), its performance is much more sensitive on the quality of the MSA used as input than methods based on machine learning. Thus, GEMME strikes us as an optimal proxy for whether or not resource-saving alignment methods such as MMseqs2 suffice for variant effect prediction. We placed ourselves in a context where GEMME relied solely on the information contained in a single input MSA to make the predictions (**Figure 1**). This setup allows for a fair comparison of different MSA generation protocols. It contrasts with the original publication (Laine et al., 2019) where GEMME would exploit two sets of input sequences. We assessed GEMME predictions against a large collection of 87 Deep Mutational Scanning experiments (DMS) totalling *∼*1.5M missense variants across 72 diverse protein families (Notin et al., 2022) (**Additional file 1: Figure S1**). We used the Spearman rank correlation coefficient to quantify the accuracy of the predictions, as previously done by us and others (Notin et al., 2022; Meier et al., 2021; Laine et al., 2019).

We show that the expand-and-filter many-to-many sequence search strategy implemented in ColabFold yields the highest-quality mutational landscapes for most of the proteins. For edge cases, where the filter is too drastic, we propose a simple solution to overcome the issue. We facilitated the import of alignments generated by ColabFold into the GEMME webserver, simplifying accessibility for users at: http://www.lcqb.upmc.fr/GEMME. Moreover, we provide predictions for the entire human proteome at: https://doi.org/10.5061/dryad.vdncjsz1s. The other datasets generated and/or analysed during the current study are available in the same Dryad repository.

## Results and Discussion

We refer to the four different MSA generation protocols and resources we considered as ColabFold, ProteinGym-MSA, ProteinNet and Pfam (see *Methods*, **Figure 1**, and **Additional file 1: Table S1**). They all proved useful for several applications, and they represent a variety of choices in terms of sequence database, search algorithm and sequence context. In short, ProteinGym-MSA relies on the profile HMM-based method JackHMMer (Eddy, 2011) to search sequences against UniRef100 (Suzek et al., 2015), a non-redundant version of UniProt (uni, 2023). The MSAs generated with this protocol have been widely used to assess mutational outcome predictors (Notin et al., 2022; Hopf et al., 2017). ColabFold uses the many-against-many sequence search algorithm MMseqs2 against the same database as ProteinGym, namely UniRef100. The MMseqs2 search strategy differs markedly from JackHMMer in that it uses the 30% sequence identity clustered database UniRef30 (Mirdita et al., 2022) as a proxy to Uniref100 to select sequences. This strategy involves a series of expansion and filtering steps with different thresholds for which straightforward equivalents are not available in JackHMMer. Furthermore, ColabFold offers the possibility to include metagenomic data from the Big Fantastic Database (BFD) (Jumper et al., 2021). Both ProteinNet and Pfam are large readily available resources of MSAs generated from profile HMMs. Their advantage compared to the two other protocols is that they do not add any computational overhead on top of GEMME prediction itself.

One potential drawback though is that they typically do not cover the full protein length and thus lack contextual information. Specifically, ProteinNet focuses on protein regions whose 3D structures have been experimentally resolved. It uses JackHMMer against Uniprot Archive (Uniparc) (Consortium et al., 2018) and a collection of metagenomic sequences (Nordberg et al., 2014). Pfam is centered on manually curated protein domains, and we used the largest available MSAs, generated with HMMer against the whole UniProtKB. We chose to adopt the default parameters settings for each considered protocol or resource. This choice guarantees that our findings are comparable to those reported in the literature for these resources and that users can reproduce our results without fine-tuning the parameters or algorithms.

### ColabFold alignments yield high-quality mutational landscapes with fewer sequences

We found that ColabFold and ProteinGym-MSA were the best performing protocols and the only ones covering all *∼*1.5M mutations from the ProteinGym benchmark (**Table 1**). The MMseqs2-based ColabFold search strategy consistently yielded better predictions than the JackHMMer-based ProteinGym-MSA protocol for two thirds of the DMS (**Figure 2A**). This result holds true whether the ColabFold protocol was performed against the union of UniRef100 and the ColabFold environmental database, which is the default set up, or only against UniRef100, *i.e.* the same database as used by ProteinGym-MSA (**Additional file 1: Figure S2**). Moreover, the expand-and-filter strategy implemented in ColabFold produced shallower alignments, with substantially fewer sequences, than the other protocols (**Additional file 1: Table S1 and Figure S3**). For instance, all proteins falling in the ’high’ alignment depth category (*N_eff_ /L >* 100, see *Methods*) based on their ProteinGym-MSA alignments, would be reclassified in the ’medium’ category (1 *< N_eff_ /L <* 100) based on their ColabFold MSAs (**Figure 2B**, red triangles, and **Additional file 1: Figure S4**). This decreased alignment depth is accompanied by an improved prediction accuracy, by an average Spearman rank correlation difference Δ*ρ̄* = 0.032, underlying the relevance of the ColabFold search strategy for these proteins. ColabFold also produced shallower alignments for most of the proteins from the ’medium’ category (**Figure 2B**, blue triangles). The differences in alignment depths have a limited impact on the prediction accuracy except for two proteins, namely the polymerases PA and PB2 from the influenza A virus (**Figure 2B**, see the two outliers). For these two extreme cases, the ColabFold MSAs are 20 times shallower than those produced by ProteinGym-MSA, resulting in a lower prediction accuracy by Δ*ρ ∼ −*0.3. The reason behind such a difference is the low divergence of these protein families. Indeed, the ProteinGym-MSA alignments contain a few tens of thousands of sequences, but almost all of them are very similar to the query (**Additional file 1: Figure S5A-B**, middle panels). GEMME is still able to exploit this limited variability to produce good-quality predictions (*ρ* values of 0.586 and 0.435). However, ColabFold’s strategy massively filtered out these similar sequences, down to a few tens (**Additional file 1: Figure S5A-B**, left panels). It brought in more divergent sequences, but they did not counterbalance the loss of information and GEMME predictions dramatically deteriorated. Removing the stringent filter of ColabFold and thereby expanding the MSAs, allowed for the restoration of prediction accuracies similar to those achieved by ProteinGym-MSA (**Additional file 1: Figure S5A-B, right panels**). We further identified two other proteins from the benchmark for which the ColabFold alignments had few sequences (less than 200). We obtained a significant gain in performance by removing the filter for these two additional cases (**Additional file 1: Figure S5C-D**). Although the number of concerned proteins in the benchmark remains small, this result suggests that removing the filter when the alignment contains less than 200 sequences can be beneficial. A condition for this no-filter strategy to be effective is the presence of numerous highly similar sequences, as is often the case for viral protein families. Finally, ColabFold’s default strategy expanded the MSAs for all proteins belonging to the ’low’ category, resulting in a small gain in the overall performance (**Figure 2B**, green triangles).

**Table 1:** Average Spearman’s rank correlation between predicted values and experimental measurements on the ProteinGym substitution benchmark. The *N_eff_* categories *Low*, *Medium* and *High* were taken from (Notin et al., 2022) and correspond to the ProteinGym-MSA alignments. We use this classification as a reference, although proteins may change category between the different protocols (see Additional file 1: Figure S4). The Spearman rank correlations are computed either over all residues from the target sequences, or only the residue ranges covered by ProteinNet and Pfam, respectively. For each alignment depth category or taxon, the best performing protocol is highlighted in bold. The correlations over the full-length versus partial proteins are comparable for ColabFold and ProteinGym-MSA protocols (Additional file 1: Figure S7).

| Set | Class | #(proteins) | #(DMS) | ColabFold | ProteinGym-MSA | ProteinNet | Pfam |
| --- | --- | --- | --- | --- | --- | --- | --- |
| All |  | 72 | 87 | **0.470** | 0.463 | - | - |
|  | Low | 14 | 20 | **0.453** | 0.444 | - | - |
|  | Medium | 43 | 17 | 0.443 | **0.446** | - | - |
|  | High | 15 | 50 | **0.552** | 0.520 | - | - |
|  | Human | 26 | 32 | **0.445** | 0.436 | - | - |
|  | Eukaryote | 10 | 13 | **0.500** | 0.479 | - | - |
|  | Prokaryote | 17 | 21 | **0.529** | 0.505 | - | - |
|  | Virus | 19 | 21 | 0.429 | **0.451** | - | - |
| ProteinNet |  | 42 | 51 | **0.507** | 0.497 | 0.495 | - |
|  | Human | 19 | 23 | **0.484** | 0.466 | 0.477 | - |
|  | Eukaryote | 6 | 7 | **0.539** | 0.531 | 0.495 | - |
|  | Prokaryote | 13 | 17 | **0.562** | 0.536 | 0.540 | - |
|  | Virus | 4 | 4 | 0.353 | **0.453** | 0.410 | - |
| Pfam |  | 39 | 52 | **0.463** | 0.440 | - | 0.432 |
|  | Human | 15 | 20 | **0.440** | 0.423 | - | 0.407 |
|  | Eukaryote | 7 | 10 | **0.462** | 0.448 | - | 0.436 |
|  | Prokaryote | 9 | 13 | **0.517** | 0.489 | - | 0.496 |
|  | Virus | 8 | 9 | **0.438** | 0.399 | - | 0.391 |

**Fig. 2:**
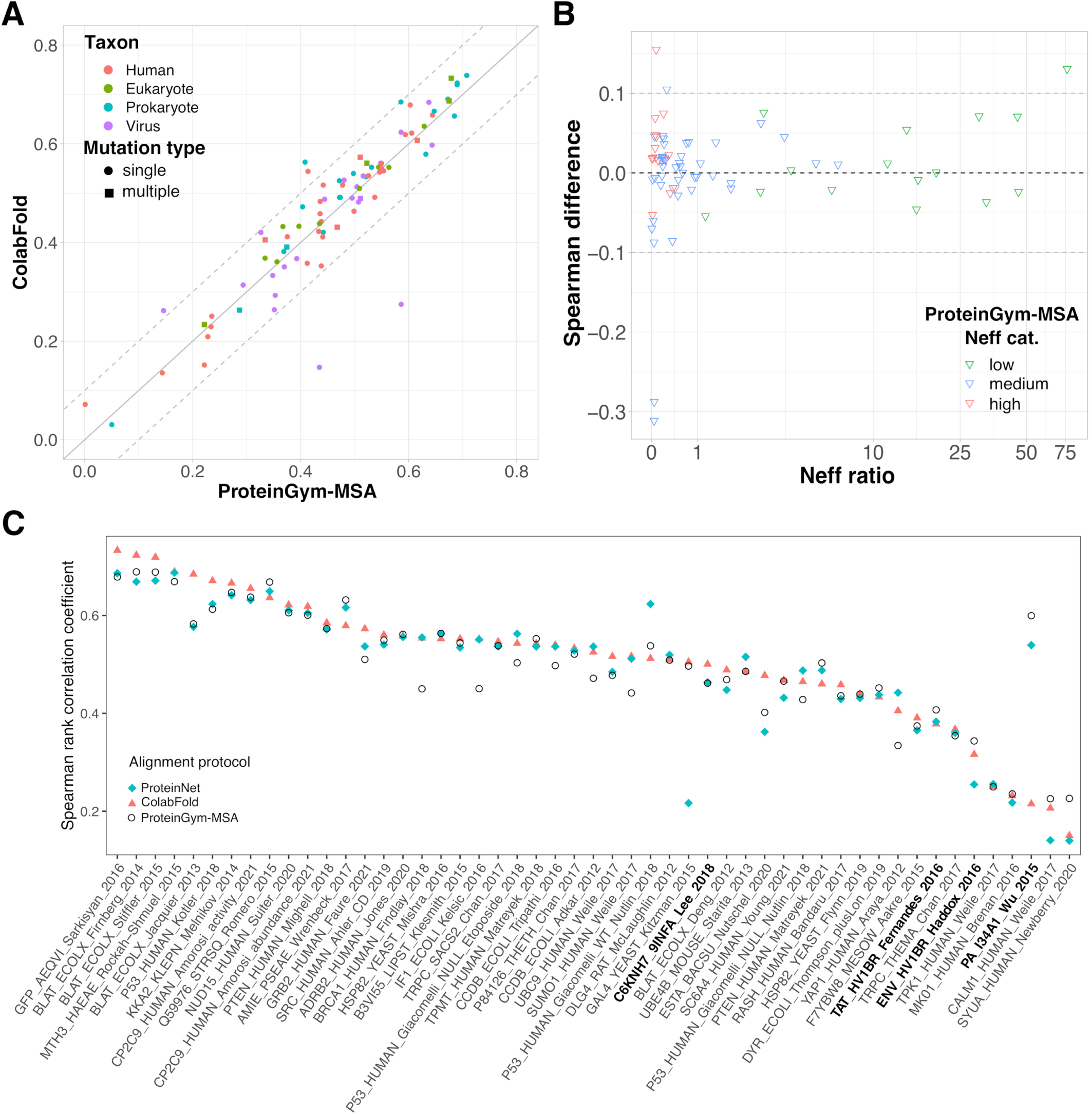
Performance comparison between the different MSA generation protocols. A. GEMME’s Spearman rank correlation coefficients (*ρ*) computed against the 87 DMS sets from the ProteinGym substitution benchmark. The input MSAs were generated using the ProteinGym-MSA (x-axis) or ColabFold (y-axis) protocols. The colors indicate the taxons of the target sequences and the shapes indicate whether the experiment contains only single mutations (circle) or also multiple mutations (square). **B.** Differences in *ρ* values in function of the number of effective sequence (*N_eff_*) ratio (**Additional file 1: Supplementary Methods**). Positive values correspond to ColabFold performing better than ProteinGym-MSA. Each point (triangle) corresponds to a given input MSA (*i.e.* a given target sequence) and its y-value is averaged over the set of DMS experiments (between 1 and 4, see **Additional file 1: Figure S1**) associated to it. The colors indicate the depth of the ProteinGym-MSA alignments, either low, medium or high, as defined in (Notin et al., 2022) (see also *Methods*). **C.** Comparison of ProteinNet, ColabFold and ProteinGym-MSA against the 51 DMS covered by ProteinNet (x-axis). The *ρ* coefficients are computed over the residue spans covered by ProteinNet alignments for all methods. The DMS associated to viral proteins are highlighted in bold.

### Environmental sequences marginally contribute to improving predictions

We assessed the contribution of the environmental sequences in the context of many-to-many sequence search with MMseqs2 and pHMMs-based search with JackHMMer (**Figure 2C** and **Additional file 1: Figure S6**). Augmenting Uniref100’s set of annotated sequences with environmental sequences expands the ColabFold MSAs by up to 3 folds without significantly impacting the mutational landscape quality of most proteins (**Additional file 1: Figure S6A**). It slightly improved prediction accuracy for the four above-mentioned viral proteins, yet without allowing reaching a good agreement with the experimental measurements – the Spearman rank correlation remains below 0.3 (**Additional file 1: Figure S6A**, see purple dots at the bottom left). By contrast, it significantly deteriorated the predictions for the human KCNH2 by Δ*ρ* = *−*0.14 (**Additional file 1: Figure S6A**, red outlier). The limited influence of metagenomics can also be observed when using JackHMMer as the search algorithm, as attested by the similar performance obtained for ProteinGym-MSA and ProteinNet (**Table 1**). Both protocols rely on JackHMMer as the search algorithm, but while ProteinGym-MSA considers only annotated sequences from UniRef100, ProteinNet searches against the UniParc archive, grouping several databases of annotated sequences, and the IMG environmental database. This expanding search results in alignments containing 3 times more sequences on average. However, we identified only a few human proteins, namely P53, BRCA1, SUMO1, and YAP1, as well as IF1 and CCDB from *E. coli*, that benefited from this additional information by up to Δ*ρ* = 0.11 (**Figure 2C** and **Additional file 1: Figure S6B**).

### Mutational landscapes of curated domains and folded regions are not better resolved

One may wonder whether the predictions are better in regions annotated as protein domains or with experimentally resolved 3D structures compared to unannotated or disordered regions. To test this hypothesis, we compared the prediction performance achieved for the full mutational landscapes versus partial landscapes focusing only on the regions covered by Pfam or ProteinNet (**Additional file 1: Figure S7**). In all cases, we used the full-length alignments generated with ColabFold or ProteinGym-MSA and ran GEMME over the entire proteins. We focused on specific regions only for the computation of the Spearman rank correlation coefficients. We did not observe any significant differences between the full-length and region-focused *ρ* distributions (**Additional file 1: Figure S7**).

Full-length alignments may display unbalanced depths over the different domains of a protein, potentially biasing the extraction of signals relevant to mutational outcomes. In order to assess the influence of the sequence context, we compared GEMME mutational landscapes predicted from full-length alignments with landscapes reconstructed from predictions obtained with domain-centered alignments (**Additional file 1: Figure S8**). Specifically, we ran GEMME on each of the Pfam alignments associated to a given protein, each one representing a curated Pfam domain, and we merged the predictions in a single landscape. We observed that the landscapes derived from full-length alignments were consistely more accurate than the reconstructed ones (**Additional file 1: Figure S8**). Indeed, the ColabFold strategy led to a higher Spearman rank correlation than the Pfam protocol for 70% of the considered DMS (**Additional file 1: Figure S9**). For the remaining 30%, the gain brought by Pfam does not exceed Δ*ρ_max_* = 0.077. Along this line, the yeast protein GAL4 gives an illustration of the importance of the extent of the sequence context (**Figure 2C** and **Additional file 1: Figure S10**). While the ProteinGym-MSA protocol could retrieve 16,159 sequences by querying the full-length query, the ProteinNet protocol retrieved only 249 sequences by querying a very small portion of the protein (6% that is 55 residues out of 881, PDB code: 1HBW). As a consequence, ProteinNet yielded a mutational landscape of a much poorer quality compared to ProteinGym, with a Spearman rank correlation of 0.217 versus 0.497 computed over the same residue range.

Expanding on our assessment against the ProteinGym benchmark, we scaled the application of GEMME using ColabFold alignments to the entire human proteome. GEMME produced predictions for 20 339 proteins (out of a total of 20 484, see *Materials and Methods*) ranging from 21 to 14 507 residues (**Additional file 1: Figure S11**). It computed all mutational landscapes exploiting the full sequence context of each protein.

## Conclusion

Multiple sequence alignments are critical to many protein-related questions. For instance, the last edition of the Critical Assessment of Structure Prediction (CASP, round 15) showed that MSA-based methods still significantly outperform protein language models in predicting protein 3D structures (Rigden et al., 2023; Elofsson, 2023). In this report, we assessed the influence of the search algorithm and the database choice for generating MSAs on the quality of *in silico* protein mutational landscapes. We ensured a clear readout of the input alignments using an unsupervised predictor relying on a few biologically meaningful parameters. The MMseqs2-based strategy implemented in ColabFold showed a good balance between prediction accuracy and computational time. It yields the best overall performance on a set of 87 DMS spanning a wide variety of proteins and covers protein regions lacking structural data or domain annotations. By controlling the number of sequences, it allows running these algorithms on machines with less memory. It is faster than classical homology detection methods by orders of magnitude. The users can easily tune the parameters, *e.g.* relax the filtering criteria, for handling protein families with low divergence. We also showed that readily available resources such as ProteinNet and Pfam are valid options, albeit only partially covering the query proteins.

In the last couple of years, a lot of attention has been drawn to optimizing, ensembling, clustering, subsampling, and pairing alignments toward improving protein 3D models (Petti et al., 2023), generating multiple functional conformations (Wayment-Steele et al., 2022), and resolving interactomes (Bret et al., 2023; Bryant et al., 2022). In the context of disease variants calling, Jagota and co-authors recently showed that vertebrate alignments exhibit a strong signal that can be used to boost specificity (Jagota et al., 2022). Nevertheless, determining which alignments are the most suitable for a given task, predictive method, or biological system often remains challenging. Our findings demonstrated that the alignment depth is not as good an indicator of prediction accuracy as one might expect. Shallow alignments can yield Spearman rank correlation as high as 0.7, and above a certain threshold, adding more sequences does not improve the predictions. Achieving accurate predictions with shallower alignments makes it possible to shed light on the mutational landscapes of protein families with few members or low divergence and also significantly reduces computational burden. In addition, we observed that extending the sequence search space to environmental datasets only marginally improves the accuracy of the predictions. Finally, we found that it is beneficial to make predictions with the knowledge of the full sequence context, rather than focusing on individual domains and concatenating the predictions afterwards. This result emphasises the importance of long-range inter-residue dependencies and suggests that deep learning methods are strongly limited by the maximal input sequence length, and thus context, they can handle.

By establishing that fast MSA generation by MMseqs2 suffices, this study demonstrates the feasibility of MSA-based computational scans of entire proteomes at a very large scale. Combining ColabFold with GEMME, it takes only a few days to generate the complete single-mutational landscape of the human proteome on the supercomputer “MeSU” of Sorbonne University (64 CPUs from Intel Xeon E5-4650L processors, 910GB shared RAM memory). We made our human proteome-scale predictions available to the community. Moreover, our findings imply ways to save resources for other MSA-based methods.

## Methods

### DMS benchmark set

We downloaded the ProteinGym substitution benchmark (Notin et al., 2022) from the following repository: https://github.com/OATML-Markslab/Tranception. It contains measurements from 87 DMS collected for 72 proteins of various sizes (between 72 and 3,423 residue long), functions (*e.g.* kinases, ion channels, g-protein coupled receptors, polymerases, transcription factors, tumor suppressors), and origins (**Additional file 1: Figure S1A-C**). The DMS cover a wide range of functional properties, including thermostability, ligand binding, aggregation, viral replication, and drug resistance. Up to four experiments are reported for each protein (**Additional file 1: Figure S1D**). Although the benchmark mostly focuses on single point mutations, it also reports multiple amino-acid variant measurements for 11 proteins (**Additional file 2: Table S2**). In the following, we considered the whole benchmark, and also a non-redundant version comprising only 59 proteins. We extracted these proteins with an adjusted version of UniqueProt (https://rostlab.org/owiki/index.php/Uniqueprot) (Olenyi et al., 2022; Mika and Rost, 2003). Compared to the original UniqueProt protocol, we used MMseqs2 instead of PSI-BLAST to improve runtime, and discarded alignments of less than 50 residues for pairs of sequences with at least 180 residues to prevent very short alignments from removing longer sequences.

### MSA resources and protocols

Two protocols, ColabFold and ProteinGym-MSA, were available for all 87 DMS (from 72 proteins) from the ProteinGym benchmark. ProteinNet was available for 51 (from 42 proteins), Pfam for 52 (from 39 proteins). When comparing two methods, we reduced the Spearman rank calculations to their common positions.

**The ColabFold protocol (Mirdita et al., 2022)** relies on the very fast MMseqs2 method (Steinegger and Söding, 2017) (3 iterations) to search against UniRef100 (Suzek et al., 2015), the non-redundant version of UniProt (uni, 2023), through a 30% sequence identity clustered version (UniRef30), and a novel database compiling several environmental sequence sets (**Additional file 1: Table S1**). It maximises diversity while limiting the number of sequences through an expand-and-filter strategy. Specifically, it iteratively identifies representative hits, expand them with their cluster members, and filters the latter before adding them to the MSA. We used the same sequence queries as those defined in ProteinGym-MSA. For all but 5 proteins, the query corresponds to the full-length UniProt sequence. For each query, we generated two MSAs by searching against UniRef30 and ColabFold environmental database, respectively, and we then concatenated them.

**The ProteinGym-MSA protocol (Notin et al., 2022)** relies on the highly sensitive homology detection method JackHMMer (Eddy, 2011) (5 iterations) to search against UniRef100 (Suzek et al., 2015), the non-redundant version of UniProt (**Additional file 1: Table S1**). JackHMMer is part of the HMMer suite and is based on profile hidden Markov models (HMMs). This protocol is relatively costly, with up to several hours for a single input MSA. It was initially described in (Hopf et al., 2017) where it was designed and tested on a subset of the current ProteinGym substitution benchmark. Hence, the proteins and DMS included in ProteinGym after this seminal publication can be considered as an independent test set. The protocol proved useful for large-scale applications (Frazer et al., 2021). In this work, we took the alignments provided with the ProteinGym benchmark (Notin et al., 2022).

**The ProteinNet protocol (AlQuraishi, 2019)** also performs 5 iterations of JackHMMER, but it extends the sequence database to the whole UniProt Archive (Uniparc) (Consortium et al., 2018) complemented with metagenomic sequences from IMG (Nordberg et al., 2014) (**Additional file 1: Table S1**). Another difference from ProteinGym-MSA is that the queries correspond to sequences extracted from experimentally determined protein structures available in the PDB (Berman et al., 2002). The MSAs are readily available and organised in a series of data sets, each one encompassing all proteins structurally characterised prior to different editions of the Critical Assessment of protein Structure Prediction (CASP) (Kryshtafovych et al., 2021). We chose the most complete set, namely ProteinNet12. It covers all proteins whose structure was deposited in the PDB before 2016, the year of CASP round XII (Moult et al., 2018).

For each protein from the ProteinGym benchmark, we retrieved the corresponding PDB codes from the Uniprot website (https://www.uniprot.org) and picked up the structure with the highest coverage among those represented in ProteinNet12 (**Additional file 2: Table S2**). We could treat 42 proteins, out of 72 in total. For the remaining ones, the positions covered by the available MSAs were out of the range of mutated positions.

**The Pfam database (Mistry et al., 2021)** is a resource of manually curated protein domain families. Each family, sometimes referred to as a Pfam-A entry, is associated with a profile HMM built using a small number of representative sequences, and several MSAs. We chose to work with the full UniProt alignment, obtained by searching the family-specific profile-HMM against UniProtKB (**Additional file 1: Table S1**). The proteins sharing the same domain composition will have exactly the same MSAs. To avoid such redundancy, we focused on the non-redundant set of 59 proteins from ProteinGym. For each protein, we first retrieved its Pfam domain composition and downloaded the corresponding MSAs from the Pfam website (https://pfam.xfam.org, release 34.0). We could retrieve at least one (and up to 5) MSA overlapping with the range of mutated positions for 39 proteins (**Additional file 2: Table S2**). Each detected Pfam domain appears only once in the set.

### Alignment depth

We measured the alignment depth as the ratio of the effective number of sequences *N_eff_* by the number of positions *L*. The effective number of sequences is computed as a sum of weights (Ekeberg et al., 2013),

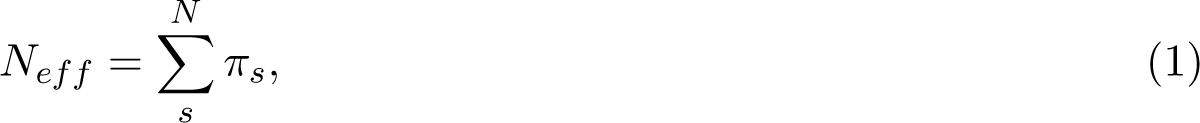

where *N* is the number of sequences in the MSA and *π_s_* is the weight assigned to sequence **x**^(^*^s^*^)^, computed as

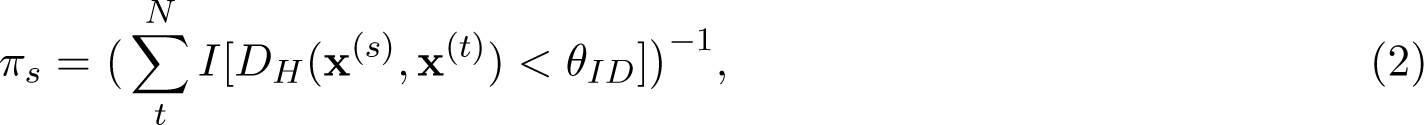

where *D_H_*(**x**^(^*^s^*^)^, **x**^(^*^t^*^)^) is the normalised Hamming distance between the sequences **x**^(^*^s^*^)^ and **x**^(^*^t^*^)^ and *θ_ID_* is a predefined neighbourhood size (percent divergence). Hence, the weight of a given sequence reflects how dissimilar it is to the other sequences in the MSA. To be consistent with (Notin et al., 2022), we set *θ_ID_* = 0.2 (80% sequence identity) for eukaryotic and prokaryotic proteins, and *θ_ID_* = 0.01 (99% sequence identity) for viral proteins.

In (Notin et al., 2022), MSAs are labeled as low, medium or high depending on the ratio *N_eff_ /L_cov_*, where *L_cov_* is the number of positions with less than 30% gaps. Specifically, MSAs with *N_eff_ /L_cov_ <* 1 are considered as shallow (’low’ group) whereas those with *N_eff_ /L_cov_ >* 100 are considered as deep (’High’ group). MSAs with 1 *< N_eff_ /L_cov_ <* 100 are in the intermediate ’Medium’ group. In our calculations, we consider the ratio between *N_eff_* and the total number of positions *L*, which is equal to the length of the target sequence for both ProteinGym-MSA and ColabFold MSAs.

### GEMME methodology and usage

GEMME exploits the evolutionary history relating the natural protein sequences to estimate the functional impact of mutations. It relies on a measure of evolutionary conservation explicitly accounting for the way protein sites are segregated along the topology of evolutionary trees (Engelen et al., 2009). A conserved position is associated with at least two subtrees of ancient origin and homogeneous with respect to that position (all sequences in a subtree display the same amino acid). Since the trees are built from global similarities between sequences, the whole sequence context plays a role in estimating the conservation level of a given position. The GEMME algorithm makes use of these conservation levels in two main steps. First, to compare different substitutions occurring at the same position, it combines amino acid frequencies, computed with a reduced alphabet, with evolutionary distances representing the mimimum amount of changes necessary to accommodate the mutations of interest. We determine the evolutionary distance associated to the substitution of *a* into *b* at position *i* as the minimal conservation-weighted Hamming distance between the query wild-type sequence and any sequence from the input alignment displaying *b* at position *i*. Then, to be able to compare substitutions occurring at different positions, GEMME weights the predicted effects with the conservation levels.

In the original GEMME publication (Laine et al., 2019), we gave two sets of sequences as input to GEMME. We used the ProteinGym-MSA protocol to generate an input alignment and we compiled an additional set of input sequences using PSI-BLAST (Altschul et al., 1997) against the NCBI’s non-redundant (NR) database (O’Leary et al., 2015) (**Figure 1**). GEMME used the later to estimate the conservation levels, and the former to computed amino acid frequencies and evolutionary distances. Since then, we observed that the additional set of sequences had a limited impact on the performance (average Δ*ρ̄* = 0.012 on the dataset reported (Hopf et al., 2017)). Hence, in more recent studies (Tsuboyama et al., 2023; Mohseni Behbahani et al., 2023), we solely relied on an input alignment generated with the ProteinGym-MSA protocol. In the present work, for all calculations, we asked GEMME to exploit only a single input MSA generated by one of the four tested protocols and resources (see **Additional file 1: Supplementary Methods** for computational details).

### Application to the human proteome

We retrieved 20 586 protein identifiers and their amino acid sequences from the Swiss-Prot reviewed human proteome available in UniProt (uni, 2023), as of August 2023. We generated MSAs with the ColabFold protocol against UniRef30 v2302 and ColabFold Environmental Database v202108. We systematically regenerated the MSAs containing less than 200 sequences without the filter step. We modified the sequences that contained undefined residues (’X’ or ’U’ symbol) in the following way. When the undefined residue was located at the beginning of the sequence, the corresponding column in the alignment was always filled with gaps, and thus we removed that column. Otherwise, we replaced the undefined residue(s) by the most frequent amino acid appearing at the corresponding position(s) in the MSA. We ran GEMME through the Docker image available at: https://hub.docker.com/r/elodielaine/gemme with default parameters. A subset of 102 sequences were too short (*≤*20 residues) to be considered as proteins and were thus not treated. Another subset of 145 proteins displayed MSAs too shallow for GEMME to estimate conservation levels. In total, GEMME generated mutational landscapes for 25 339 proteins.

## Data availability

The data underlying this article are available in the Dryad repository https://doi.org/10.5061/dryad. vdncjsz1s.

## Table and figure legends

