## Additional File 1 for "Alignment-based protein mutational landscape prediction: doing more with less"

### Supplemental methods

#### Generating the predictions with GEMME

GEMME takes as input a FASTA-formatted MSA, with the ungapped query sequence on top. We used the tool *reformat.pl* from the HH-suite [1] to convert A2M and A3M alignment files into FASTA format. For the conversion from A3M, we subsequently modified the sequence headers to keep only the first term using the *awk* command: "awk '{if(substr($0,1,1) == ">"){print $1} else {print $0}}' aliTMP.fasta > aliXXX.fasta", where *XXX* is the protein identifier. Moreover, we modified the MSAs from ProteinNet and Pfam by putting the sequence of interest on top and removing the insertions with respect to this sequence. We used GEMME's Docker image, available from `http://www.lcqb.upmc.fr/GEMME`, to compute the predictions. For the proteins with only single mutations, we predicted the full mutational landscape with the command: "python2.7 $GEMME_PATH/gemme.py aliXXX.fasta -r input -f aliXXX.fasta" where *aliXXX.fasta* is the input MSA file in FASTA format. For the proteins with multiple mutations, we predicted only the effects of the mutations of interest. To do so, we passed a file specifying the list of mutations as input with the option "-m". We used the default parameters for all proteins and all input MSAs, except for the ProteinGym-MSA alignments associated with R1AB_SARS2. In that case, the diversity of the first 20,000 sequences, retained by default for estimating conservation levels, was too low. To circumvent this issue, we took the last 80,000 sequences instead by cropping the MSA and using the option "-N".

#### Assessing and comparing the predictions

Assessing and comparing the predictions obtained from the ProteinGym-MSA and ColabFold MSAs was straightforward since they cover the entire range of mutated positions and their query sequence is identical to the wild-type sequence used in the DMS. The MSAs from ProteinNet and Pfam however typically cover only a part of the mutated region and their query sequence sometimes display a few mutations with respect to the DMS wild-type sequence. To compute the Spearman correlation, we restricted ourselves to the covered positions displaying the correct wild-type amino acid. When comparing two methods, we further reduced the calculation to their common positions.

#### Supplemental tables and figures

Supplemental Table S1: **Details about the MSA generation protocols.**

| **Name** | **Databases** | **Search algorithms** | **Fine tuning** | **#(covered proteins)[a]** | **#(sequences) Min - Max** |
|---|---|---|---|---|---|
| ColabFold | UniRef100 through UniRef30 and ColabFold environmental DB[b] [3] | MMseqs2 [2] | no | 72 | 126 - 24,269 |
| ProteinGym-MSA | UniRef100 [4] | JackHMMer [5] | yes[c] | 72 | 44 - 539,868 |
| ProteinNet | UniParc[d] [6] and IMG [7] | JackHMMer [5] | no | 42 | 249 - 1,389,216 |
| Pfam | UniProtKB [8] | HMMer [5] | yes[e] | 39[f] | 134 - 283,380 |

[a]We indicate the number of proteins treated with each protocol, out of the 72 proteins comprised in the ProteinGym substitution benchmark. [b]ColabFold environmental database contains BFD [9], which includes UniProt/TrEMBL+Swissprot, Mgnify [10], MetaEuk [11], SMAG [12], TOPAZ [13], MGV [14], GPD [15], and MetaClust2 [16]. [c]For each protein, 9 MSAs were generated by exploring bit score thresholds from 0.1 to 0.9 and the MSA leading to the highest number of significant Evolutionary Couplings [17] was retained. [d]UniParc, for UniProt Archive, is a non-redundant archive of protein sequences extracted from more than 10 public databases, including UniProtKB, Ensembl [18], PDB, FlyBase [19] and WormBase [20]. [e]For each Pfam family, the profile HMM used to query UniProtKB was hand curated, and the score threshold used to select the sequences was set manually. [f]For this protocol, we considered a non-redundant subset of 59 proteins.

Supplemental Table S2: **Coverage of the ProteinGym benchmark by the tested MSA generation protocols.** For each protein, we indicate its UniProt identifier, whether it is associated with measurements for multiple mutations, and whether the mutated region is covered by each of the tested protocols. We also give the PDB code selected for ProteinNet, and the number of Pfam domains (with available MSAs) overlapping with the mutated region.

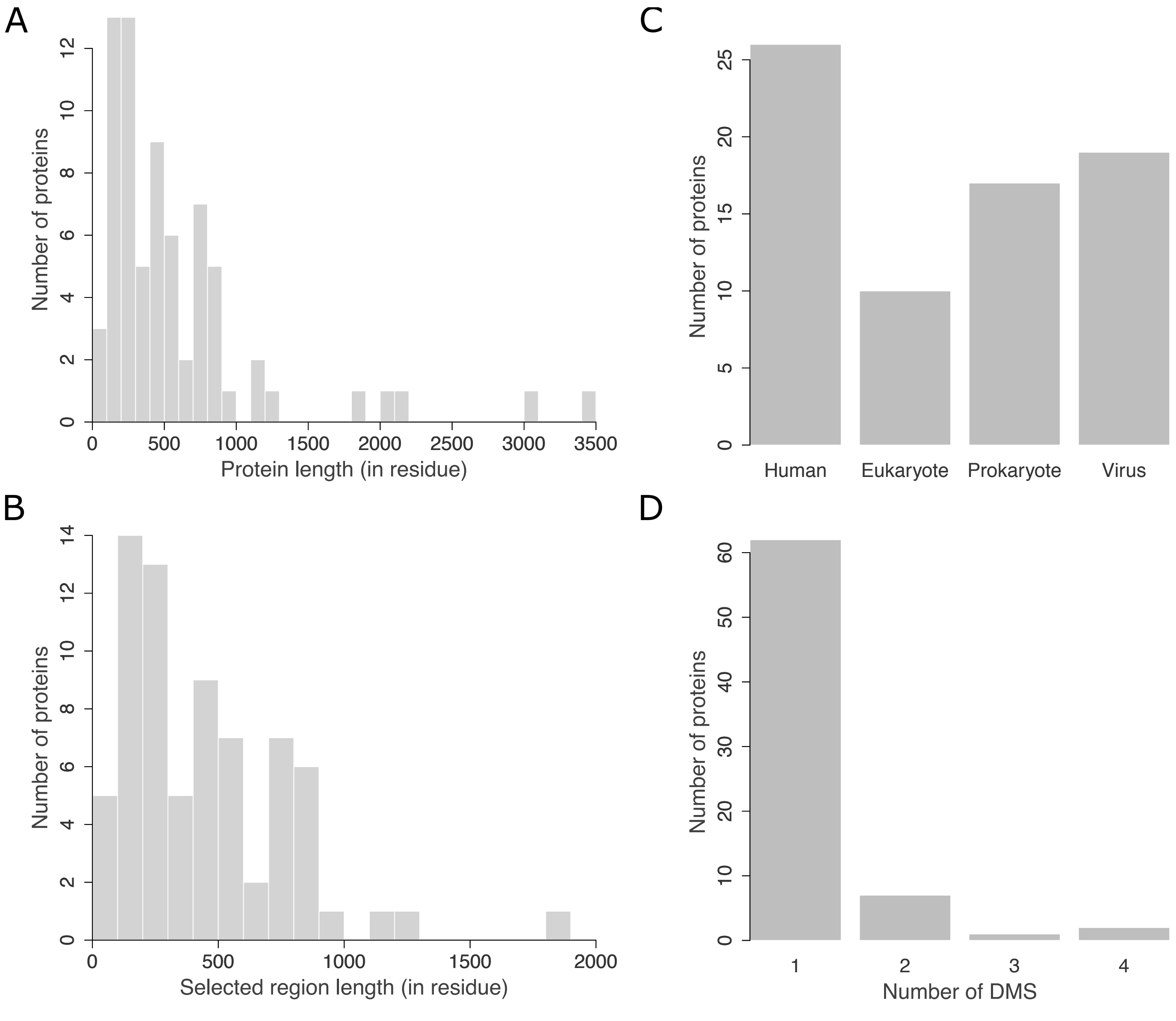


Supplemental Figure S1: **ProteinGym benchmark properties.** **A.** Distribution of the length (in number of residues) of the 73 target protein sequences from the benchmark. **B.** Distribution of the length (in number of residues) of the protein regions covered by ProteinGym-MSA alignments. **C.** Taxonomic classification of the proteins. The label "Eukaryote" refers to non-human eukaryotes. **D.** Distribution of the number of reported experiments per protein.

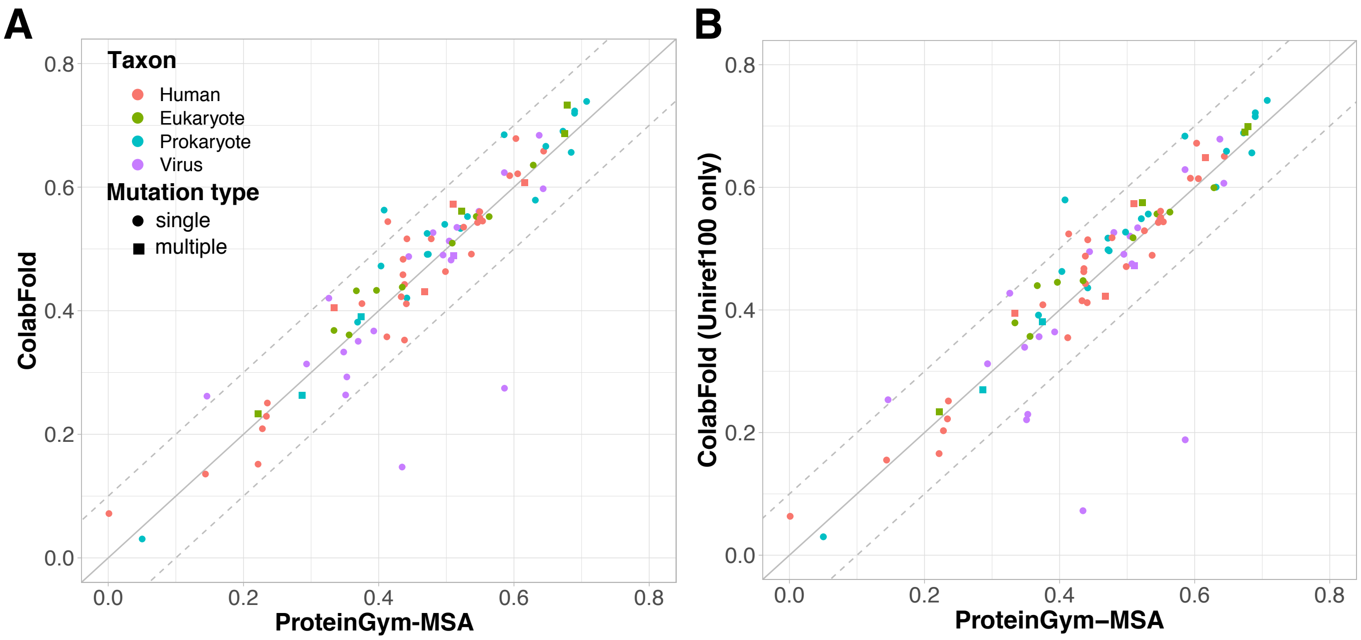


Supplemental Figure S2: **Performance comparison between ColabFold and ProteinGym-MSA.** GEMME's Spearman rank correlation coefficients ($\rho$) computed against the 87 DMS sets from the ProteinGym substitution benchmark. **A.** The input MSAs were generated using the ProteinGym-MSA (x-axis) or ColabFold (y-axis) protocols. **B.** The input MSAs come from ProteinGym-MSA (x-axis) or ColabFold's MMseqs2-based protocol against UniRef100 only (no use of the ColabFold environmental DB). The colors indicate the taxons of the target sequences and the shapes indicate whether the experiment contains only single mutations (circle) or also multiple mutations (square). Notice that panel A is identical to Figure 2A. It is reproduced here to ease visual comparison.

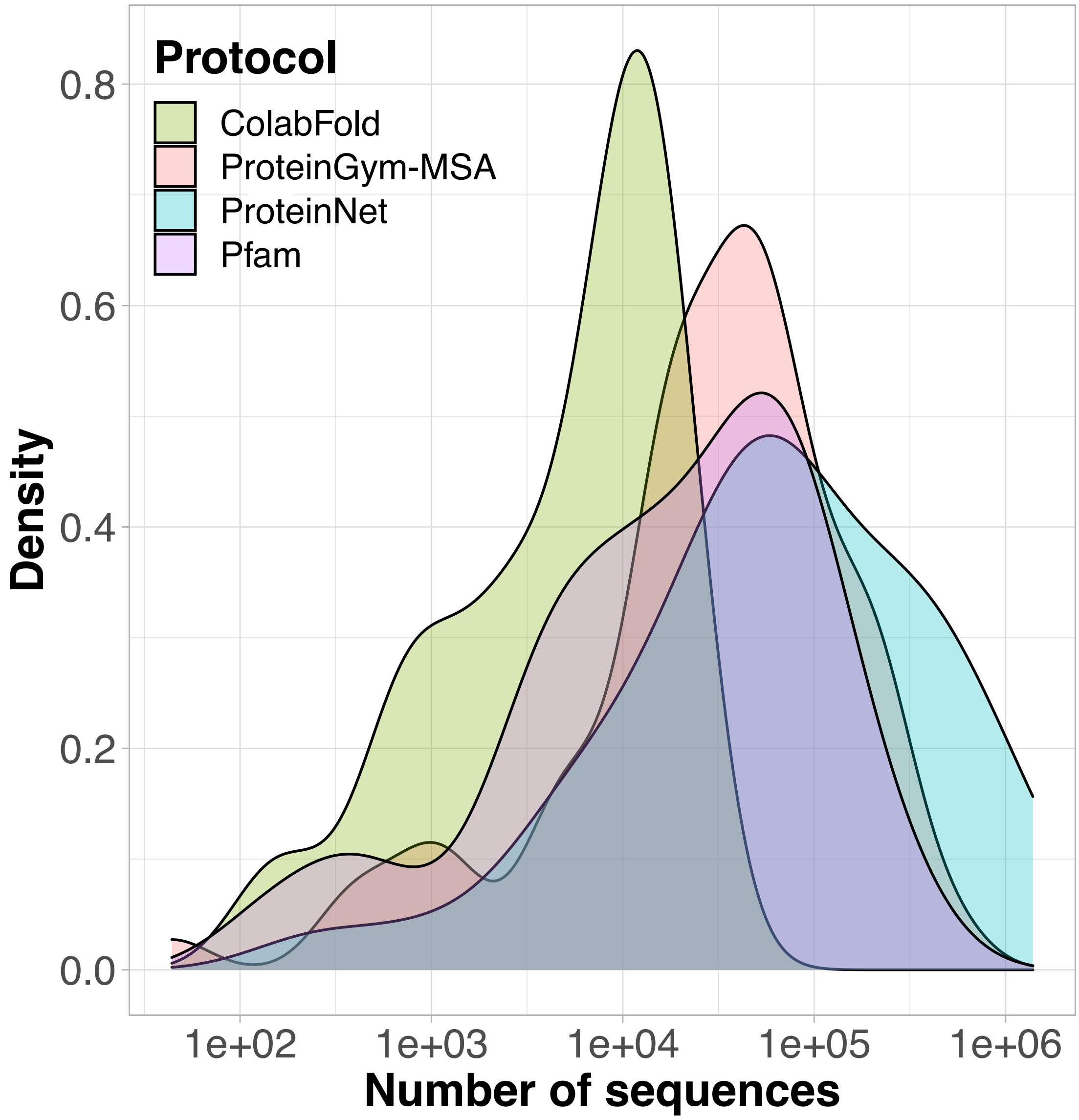


Supplemental Figure S3: **Distribution of the number of sequences per MSA depending on the protocol.** The total number of MSAs varies from one protocol to another (see full details in **Table S1**).

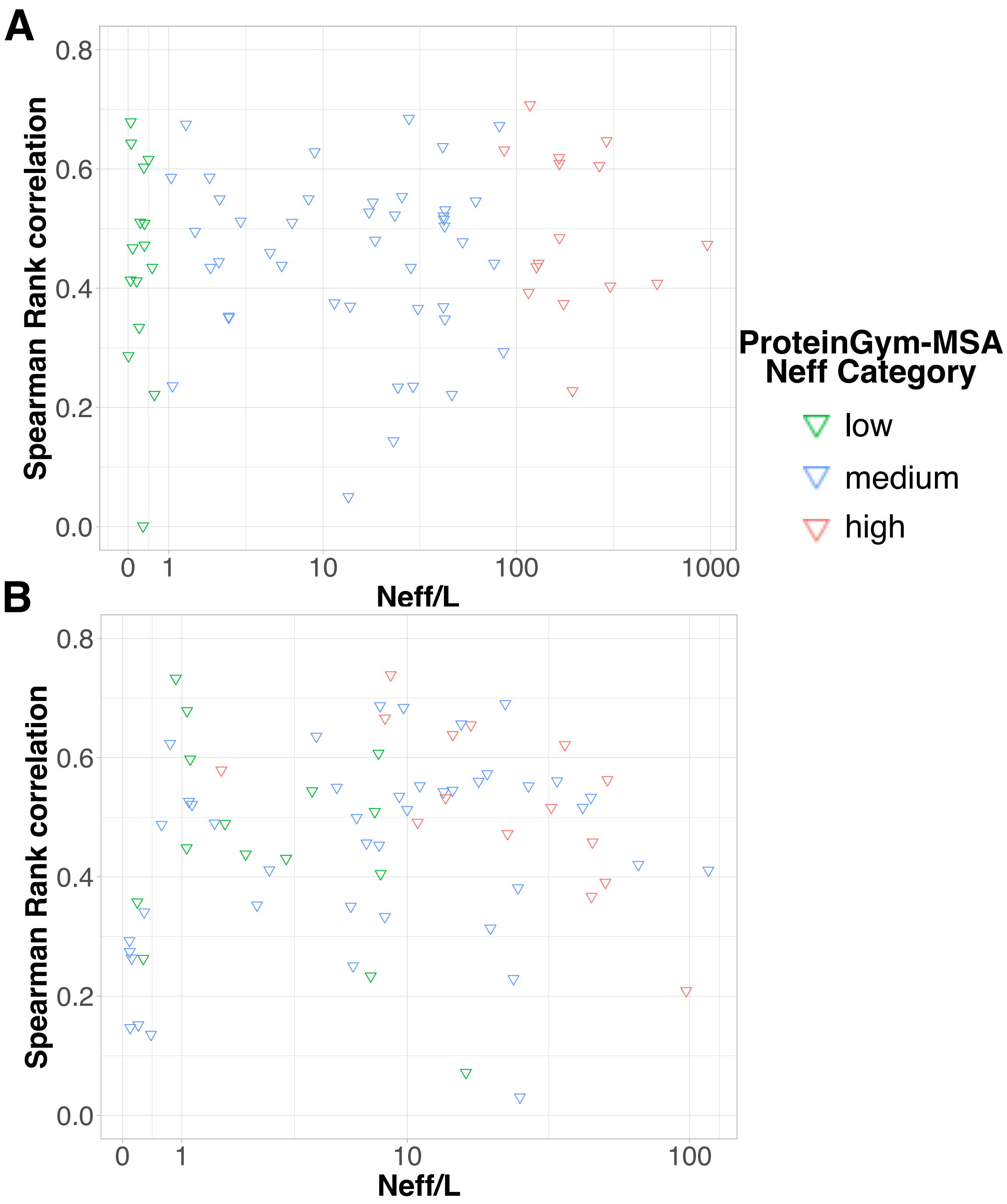


Supplemental Figure S4: **Prediction accuracy in function of the alignment depth.** The input MSAs were generated using ProteinGym-MSA (**A**) or ColabFold (**B**) protocol. Each point (triangle) corresponds to a given input MSA (*i.e.* a given target sequence) and its y-value is averaged over the set of DMS experiments (between 1 and 4, see **Fig. S1**) associated to it. The Spearman correlations computed between the y ($\rho$) and log-x ($\log N_{eff}/L$) values are 0.065 and 0.225 for ProteinGym-MSA (**A**) and ColabFold (**B**), respectively. The colors indicate the ProteinGym-MSA $N_{eff}$ categories, as defined in [21] (see also *Materials and Methods*). About half of the target sequences change category between the two protocols (see all red points, and also the blue points with a ratio lower than 1 and the green points with a ratio above 1 on panel B).

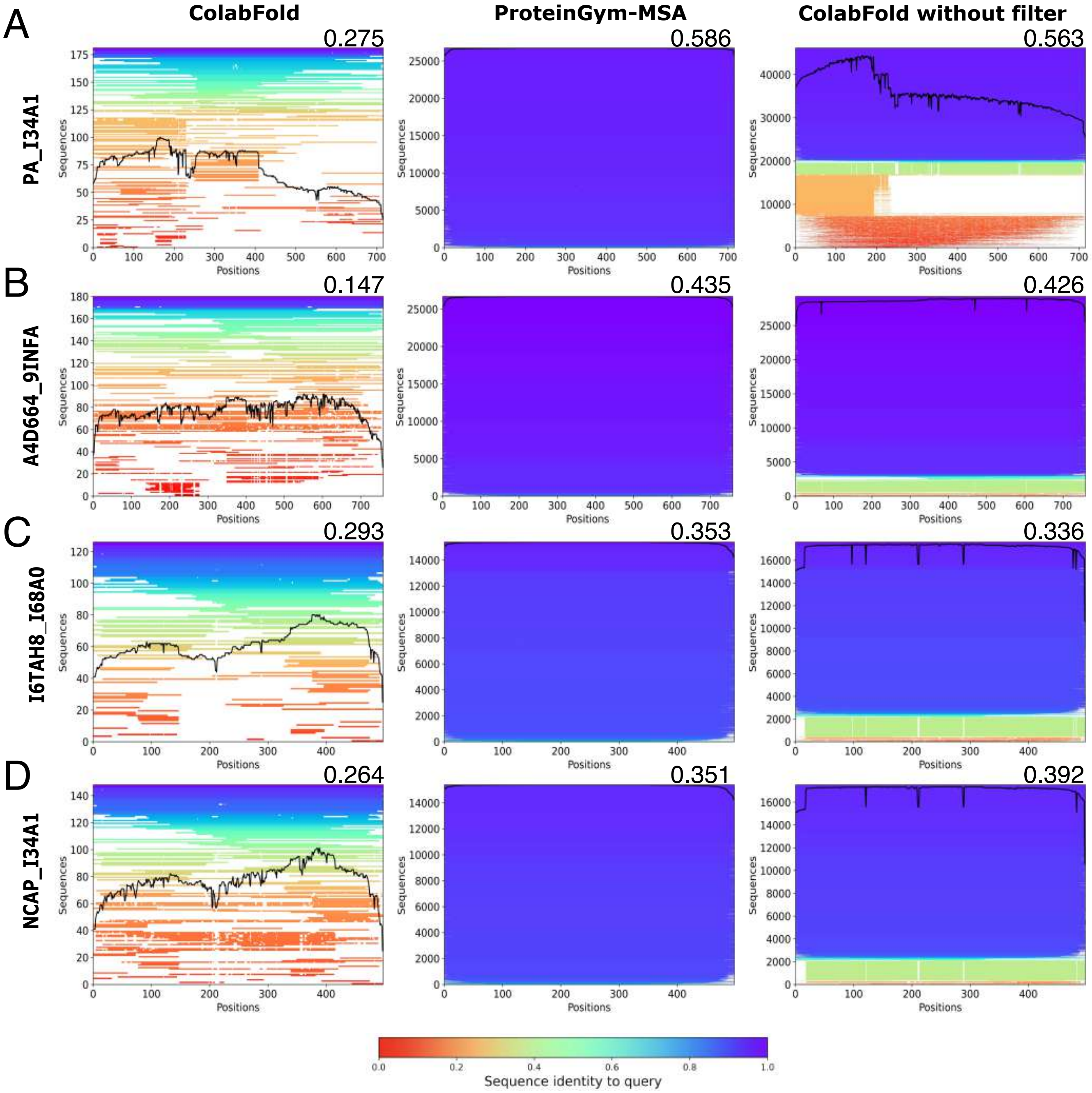


Supplemental Figure S5: **Influence of sequence filtering on GEMME performance.** Plots showing the coverage and variability of the input MSAs generated by ColabFold (left), ProteinGym-MSA (middle) and ColabFold without filter (right) for four proteins from influenza A virus, namely the polymerases PA (A) and PB2 (B), and two nucleoproteins (C-D). The corresponding Uniprot names are indicated next to the y-axes. Each line represents a sequence in the MSA and is colored according to the percentage of sequence similarity to the query (on top). Gaps are indicated by line interruptions. The black curve gives the coverage (number of non-gapped sequences) at each position. The prediction accuracy, as measured by the Spearman rank correlation coefficient, obtained from each alignment is given on the top right corner.

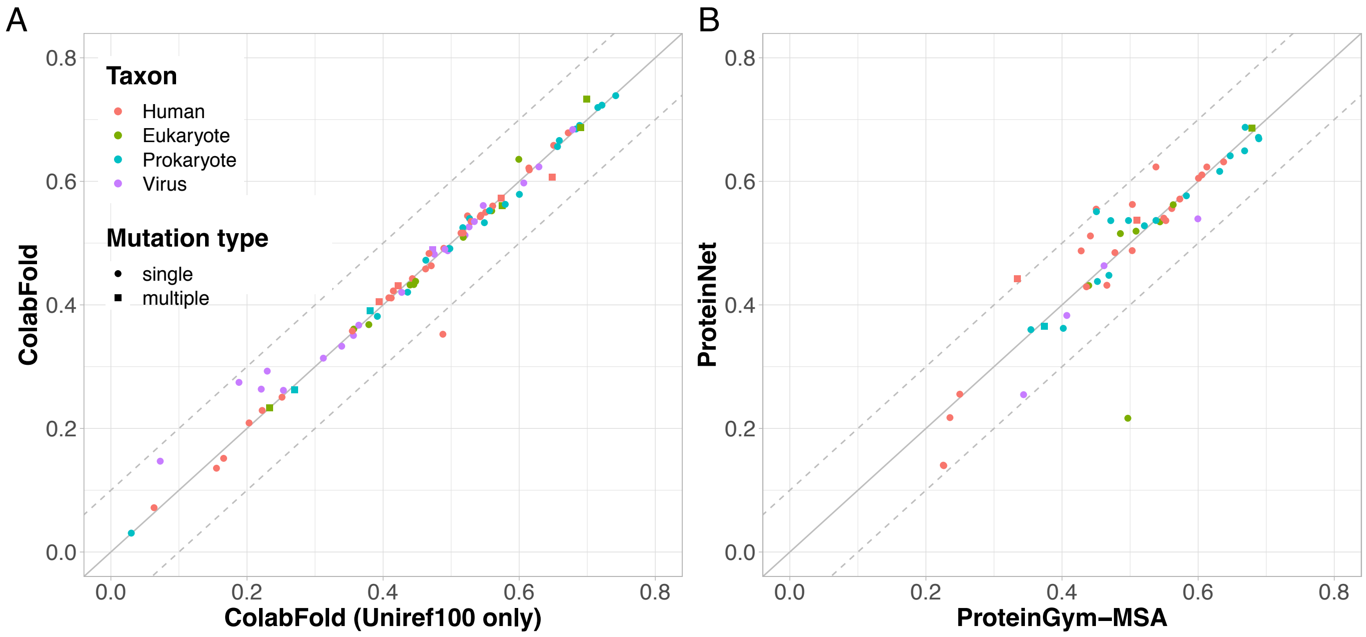


Supplemental Figure S6: **Influence of the search database on GEMME performance.** The input MSAs were generated by considering only annotated sequences (x-axis) or the union of annotated and environmental sequences (y-axis). **A.** Colabfold's MMseqs2-based many-to-many sequence search strategy. The values are reported for all 87 DMS from ProteinGym. **B.** pHMM-based search strategy implemented in JackHMMer. The values are reported for the 51 DMS covered by ProteinNet.

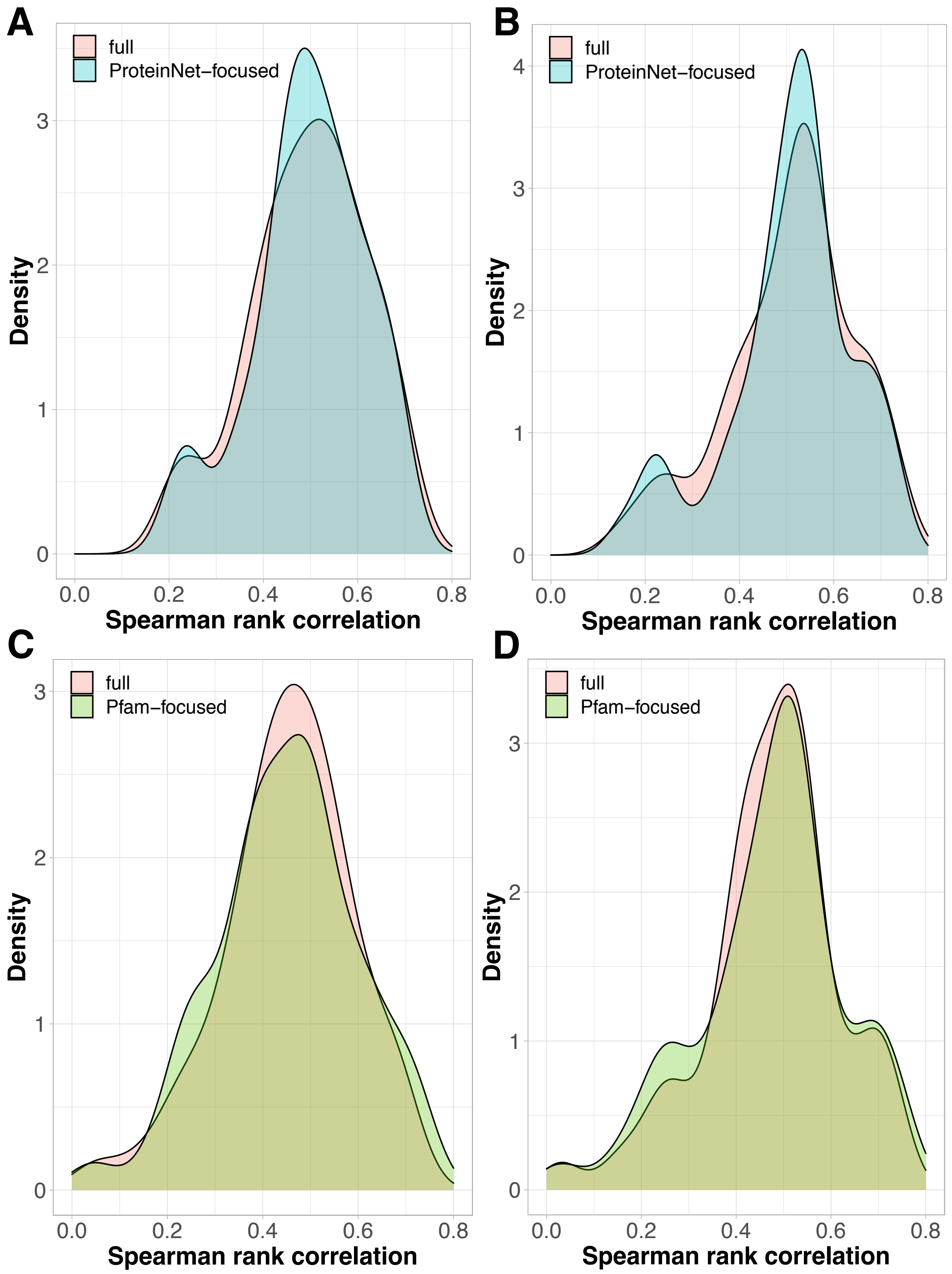


Supplemental Figure S7: **Prediction accuracy achieved on the full-length versus partial proteins.** Distributions of Spearman rank correlations obtained with the ProteinGym-MSA (A,C) and ColabFold (B,D) protocols. **A-B.** Each distribution contains 51 values corresponding to the 51 DMS covered by ProteinNet. The correlations computed over the full-length proteins (in pink) are compared to those computed over the regions covered by ProteinNet (in blue). **C-D.** Each distribution contains 52 values corresponding to the 52 DMS covered by Pfam. The correlations computed over the full-length proteins (in pink) are compared to those computed over the regions covered by Pfam (in green).

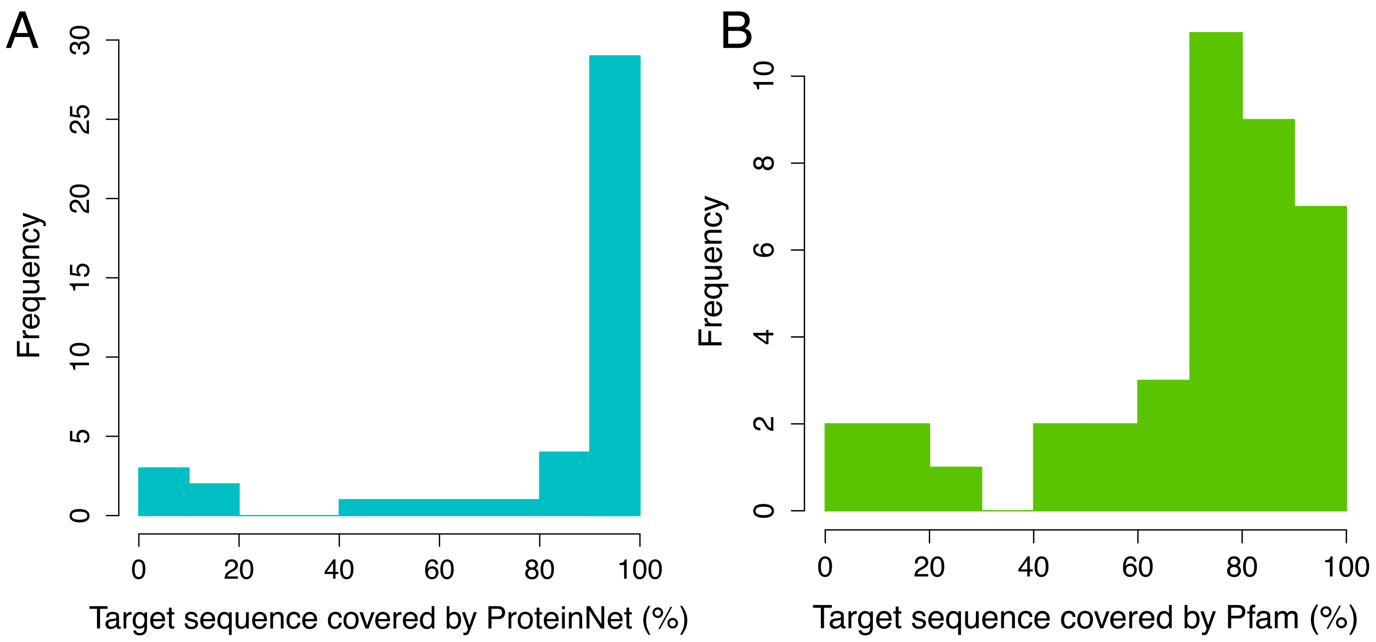


Supplemental Figure S8: **Protein coverage from the ProteinNet and Pfam MSAs.** Percentage of residues from the target sequences covered by the MSAs from ProteinNet (**A**) and Pfam (**B**).

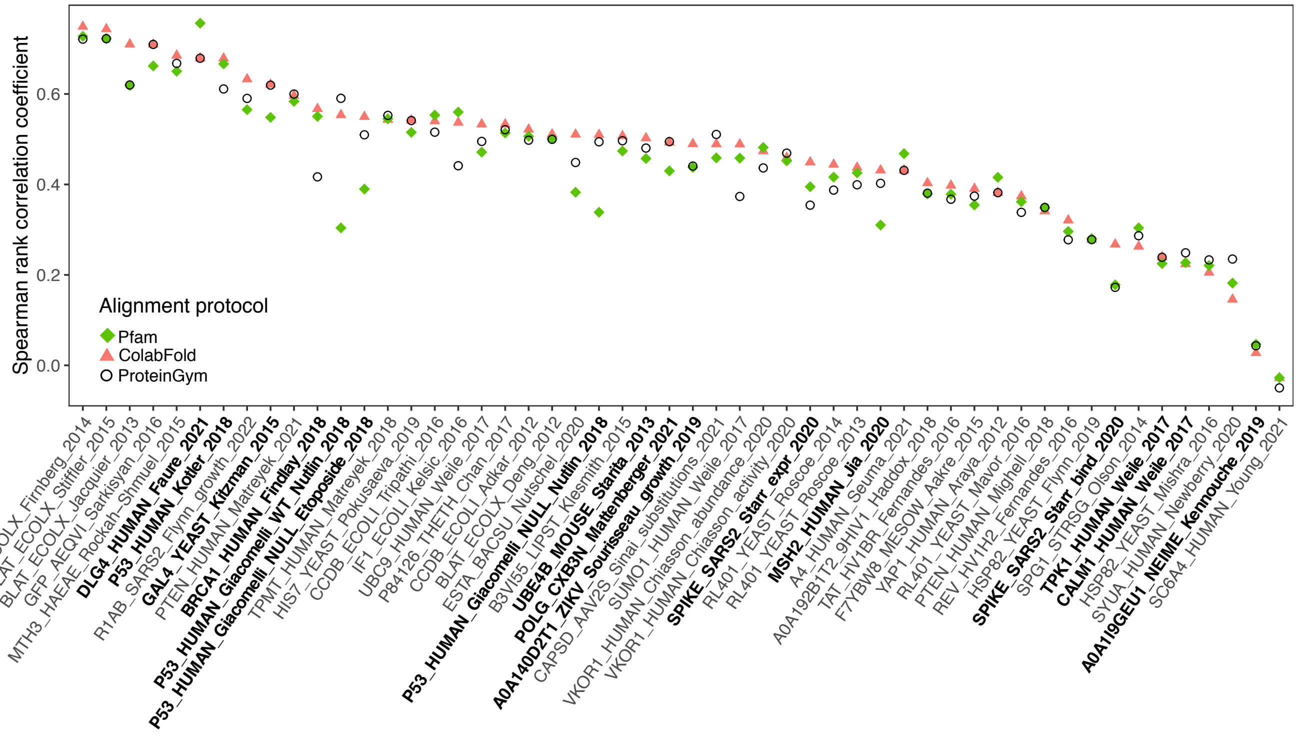


Supplemental Figure S9: **Comparison of Pfam, ColabFold and ProteinGym-MSA.** We focus on the 52 DMS covered by Pfam (x-axis). The Spearman rank correlation coefficients are computed over the residue spans covered by Pfam MSAs for all methods. The DMS associated to proteins containing more than one Pfam domains are highlighted in bold on the x-axis.

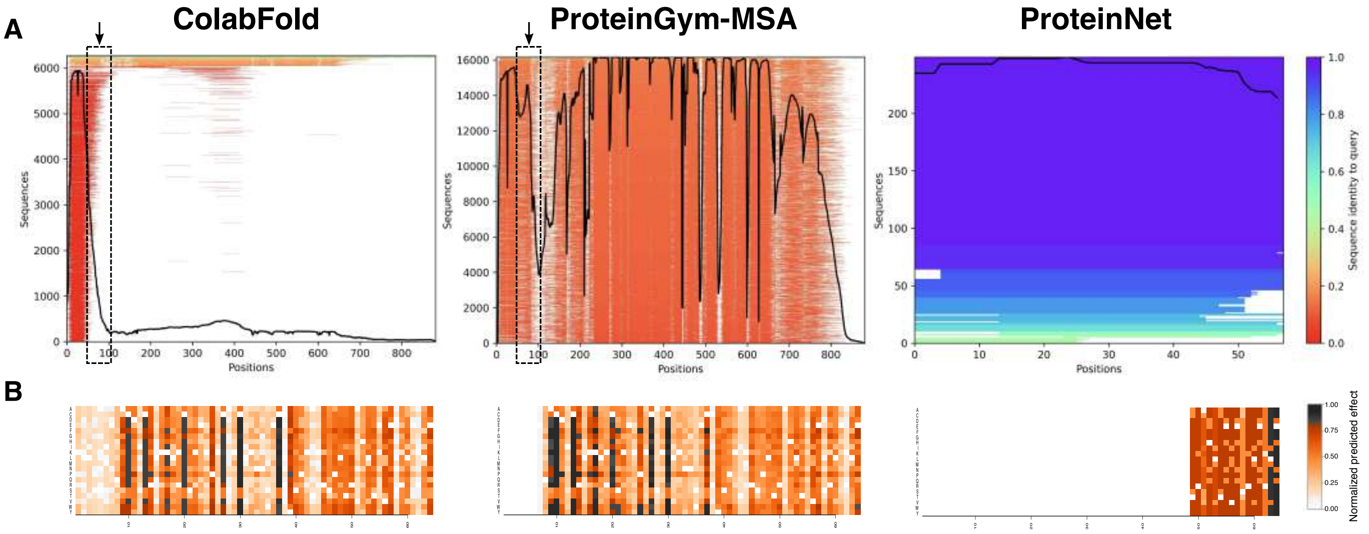


Supplemental Figure S10: **Influence of sequence context on GAL4 input MSAs and mutational landscape predictions. (A)**. Plots showing the coverage and variability of the input MSAs generated by ColabFold (left), ProteinGym-MSA (middle) and ProteinNet (right). Each line represents a sequence in the MSA and is colored according to the percentage of sequence similarity to the query (on top). Gaps are indicated by line interruptions. The black curve gives the coverage (number of non-gapped sequences) at each position. The ProteinNet alignment covers residues 50 to 106. The region is highlighted with an arrow and a dotted-line rectangle on the two other plots and we can observe that it is highly gapped. The gapped sequences were not included in the ProteinNet alignment, likely due to insufficient similarity significance. **(B)**. Mutational landscapes predicted by GEMME from the three MSAs, focusing on the region covered by the experimental DMS, namely residues 2 to 65. The darker the color, the higher the predicted mutational outcome. The values are scaled between 0 and 1 for ease of visual comparison. The white cells correspond to the wild-type amino acids.

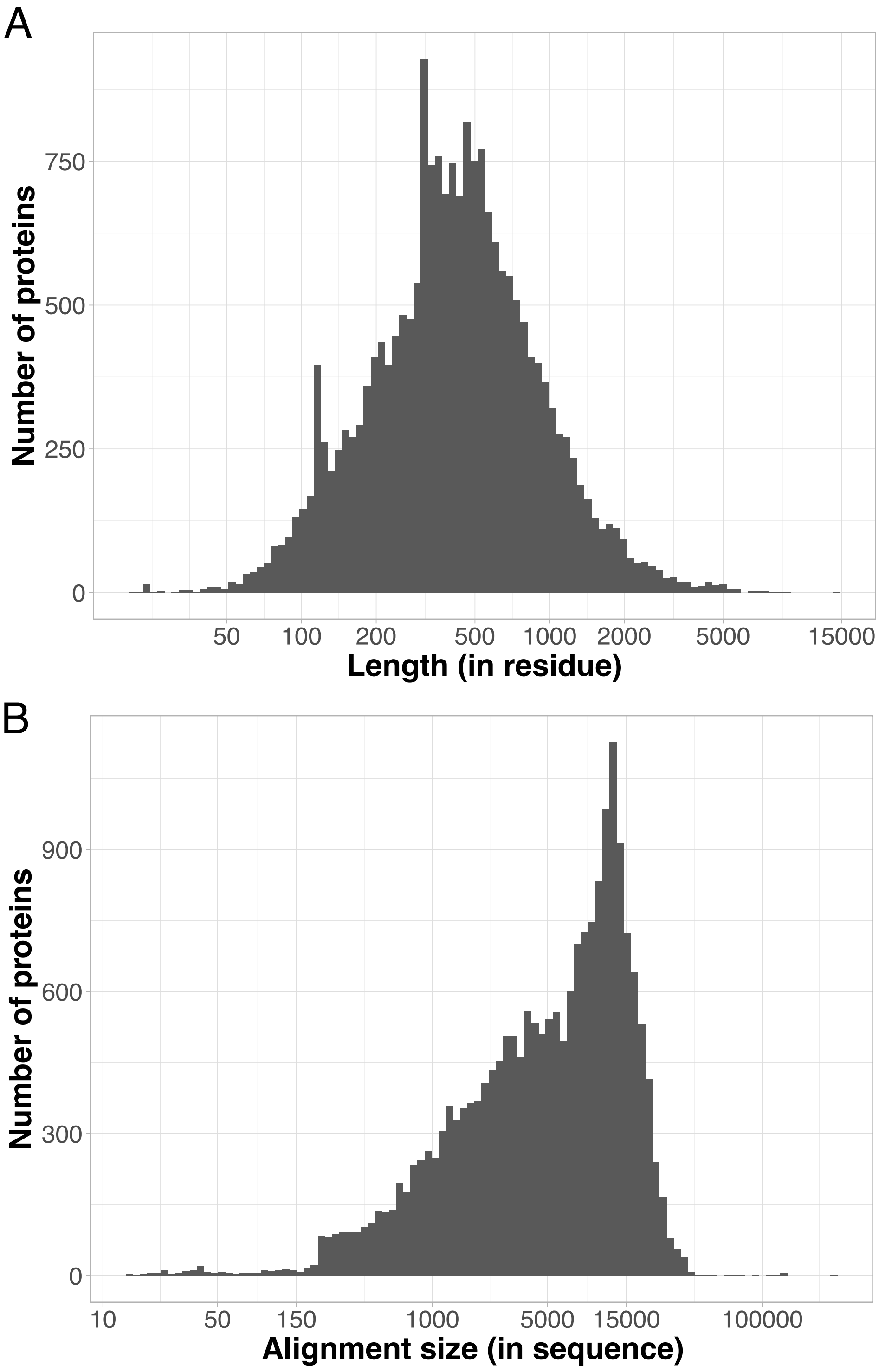


Supplemental Figure S11: **Application to the whole human proteome.** GEMME produced full-length single-mutational landscapes for 20 339 human proteins out of 20 484 (see *Materials and Methods*). **(A)**. Distribution of protein length (in residue). **(B)**. Distribution of alignment size (in sequence).

[7] Henrik Nordberg, Michael Cantor, Serge Dusheyko, Susan Hua, Alexander Poliakov, Igor Shabalov, Tatyana Smirnova, Igor V Grigoriev, and Inna Dubchak. The genome portal of the department of energy joint genome institute: 2014 updates. *Nucleic acids research*, 42(D1):D26–D31, 2014.

[8] Uniprot: the universal protein knowledgebase in 2023. *Nucleic Acids Research*, 51(D1):D523–D531, 2023.
